# Relating spinal injury-induced neuropathic pain and spontaneous afferent activity to sleep and respiratory dysfunction

**DOI:** 10.1101/2021.11.15.468636

**Authors:** S Idlett-Ali, H Kloefkorn, W Goolsby, S Hochman

## Abstract

Spinal cord injury (SCI) can induce dysfunction in a multitude of neural circuits including those that lead to impaired sleep, respiratory dysfunction and neuropathic pain. We used a lower thoracic rodent contusion SCI model - known to develop mechanosensory stimulus hypersensitivity, and spontaneous activity in primary afferents that associates neuropathic pain - and paired this with new approaches that enabled chronic capture of three state sleep and respiration to characterize dysfunction and assess possible interrelations. Noncontact electric field sensors were embedded into home cages for noninvasive capture in naturally behaving mice of the temporal evolution of sleep and respiration changes for 6 weeks after SCI. Hindlimb mechanosensitivity was assessed weekly, and terminal experiments measured primary afferent spontaneous activity *in situ* from intact lumbar dorsal root ganglia (DRG). We observed that SCI led to increased spontaneous primary afferent activity (both firing rate and the number of spontaneously active DRGs) that correlated with reduced hindpaw mechanical sensitivity, increased respiratory rate variability, and increased sleep fragmentation. This is the first study to measure and link sleep dysfunction and variability in respiratory rate in a SCI model of neuropathic pain, and thereby provide broader insight into the magnitude of overall stress burden initiated by neural circuit dysfunction after SCI.

## INTRODUCTION

Neuropathic pain can develop following peripheral or central nervous system injury, resulting from trauma or disease ^1^. Persistent and stimulus-independent neuropathic pain (i.e. spontaneous pain) develops in more than half of spinal cord injury (**SCI**) patients ^2^, yet there remains limited understanding of its underlying mechanisms ^3^ and few effective therapies for this debilitating condition.

Many studies explore neurobiological mechanisms and treatment for spontaneous pain using experimental assessment tools optimized for stimulus-evoked allodynia and hyperalgesia ^4–10^. Clinically, pain is commonly scored through questionnaires which can be highly subjective and variable ^11, 12^. Preclinically, most modalities of pain are inferred through changes in evoked mechanical or thermal sensory sensitivity, regardless of whether the pain being explored has stimulus-evoked features ^13, 14^.

Stimulus-evoked assessment methods are considered the semi-quantitative gold standard but their assessment limitations and misuse have, in part, supported arguments that spontaneous pain may represent evoked allodynia and hyperalgesia resulting from unrecognized stimuli ^3^. However, there is increasing evidence that spontaneous primary afferent activity represents one initiation site and chronic driver of maladaptive sensory processing, consistent with spontaneous pain perception ^6, 7, 15–17^. Spontaneous activity in both A and C fibers has also been observed in many neuropathic pain conditions ^7, 15, 18, 19^. In SCI, this peripheral hypersensitivity can predict the development of central pain ^20, 21^. Key studies in rodent contusion SCI models reveal potential links between stimulus-evoked hypersensitivity (hyperalgesia/allodynia) and the incidence of spontaneous firing in cell bodies of primary afferents below the level of injury ^6, 16, 22, 23^. These studies assessed spontaneous activity from C fiber dorsal root ganglia (**DRG**) neurons following DRG dissociation which may induce hyperexcitability, including increased spontaneous firing that mirrors the effects seen after chronic compression of DRGs ^24^. These limitations highlight the need to study intact DRGs to determine the presence of abnormal spontaneous DRG activity after SCI.

To better understand and measure stimulus independent spontaneous pain, non-evoked assessment methods have been developed. For example, the conditioned place preference test ^25–27^ and grimace scale ^28^ have been shown to differentiate animals with neuropathic pain from those without. Potential compounds of these tests include efficacy of analgesics, animal training, and experimenter interactions ^14, 29^. There is a need for additional indices of ongoing spontaneous pain during natural behavior with one possibility being emergent changes in other physiological phenomenon. Alternatively, home-cage wheel running has been shown to identify spontaneous migraine pain ^30^, however is inappropriate for animal models with confounding motor dysfunction such as SCI.

Impaired sleep is a significant, though often overlooked, component of suffering after SCI with meaningful impacts on life quality ^31^. Though SCI patients commonly experience reduced sleep quality ^32–35^, research on cause-effect relations between impaired sleep after SCI and additional important co-morbid conditions is limited. While it is well established that sleep disruption and chronic pain are bi-directionally and negatively reinforcing ^34–36^, their association in SCI has not been detailed clinically or in preclinical animal models. This is at least partly due to the difficulty in measuring rodent sleep. Moreover, current methods rely on invasive EEG recordings that introduce extra surgery, immune response, stress, and altered home cage environments that may complicate behavioral changes and pain after SCI.

Similarly, while there are several studies describing respiratory-related sleep dysfunction caused by cervical SCI ^34, 37, 38^, including sleep apnea ^39^, preclinical work aimed at understanding whether alterations in respiration associated with SCI -induced neuropathic pain are limited. Using lower thoracic SCI models that do not overtly impair respiratory neural circuits, studies in a restraint cylinder showed respiratory rate increases after thoracic contusion or hemisection injury that associate with the emergence of allodynia ^40, 41^. This important observation needs to be reproduced during natural behavior, which have relied on whole-body plethysmography ^42^ that has limited its applications in SCI animal models ^43^.

To better quantify spontaneous pain and address limitations of existing pain assays, we explore novel physio-behavioral indices after SCI. In the thoracic contusion SCI model of neuropathic pain, we examine metrics associated with sleep architecture and respiratory function during natural behavior in the animal’s home cage using non-invasive electric field sensors ^42, 44^. We also compare these behavioral outcomes to established mechanical sensitivity testing, along with terminal experiment observations of spontaneous activity in whole DRGs, below the level of injury.

A greater understanding of this work reveals a potential link of the pathological sensory state after SCI, with quantitative metrics of sleep and respiration abnormalities. Exploitation of this link could provide a broader perspective into understanding overall stress burden initiated by neural circuit dysfunction after SCI.

## METHODS

### Animals

All procedures were approved by the Emory University Institutional Animal Care and Use Committee. Adult C57/Bl6 mice (n = 20, female) were pair housed in home cages under standard 12:12 hour light-dark cycles with *ad libitum* food and water.

### Animal surgeries

SCI was induced via moderate contusion of the T10 spinal cord segment ^45^. Fourteen adult mice (day 205) were prepared for aseptic surgery and deeply anesthetized with 2-3% isoflurane. A midline skin incision was made to dissect the muscle and fascia prior to exposing the spinal cord at T8-T10 via a dorsal laminectomy. In six animals (Sham Group, n=6), wounds were closed at this point. In the eight remaining animals (SCI Group n=8), a moderate contusion SCI was made dorsally at T9/10 with the Infinite Horizon impactor (50 kdynes, IH-400, Precision Systems and Instrumentations) prior wound closure. All mice received post-operative pain relief (2 mg/kg meloxicam), antibiotics (2.5 mg/kg Baytril), supplemental hydration (0.5 ml sterile saline daily), and bladder expression twice daily until each animal voided independently. One animal was excluded from these data due to meeting endpoint criteria. Six additional mice served as naïve controls (Naïve, n=6).

### Physio-behavioral monitoring

Home cages (32 × 18 × 14cm, Super Mouse™ microisolator), described previously^42, 44^, were instrumented with electric field (**EF**) sensors (Plessey Semiconductors, PS25251, 1 cm^2^, +/−5V, 1 kHz sampling) able to passively translate disruptions of the local electric field caused by movement into a voltage trace. EF sensors have been validated against respective gold standard techniques to reliably quantify resting respiration, sleep-wake staging (including differentiating rapid eye movement sleep, REM, from Non-REM sleep), and animal motor activity noninvasively from outside the animals’ home cage ^42, 44^.

Animals were acclimated to their instrumented home cages for several days prior to recording (**Figure 1A**). The dividing insert was only present during physio-behavioral recordings; cage-mates shared the full home cage at all other times. Three baseline recordings were collected prior to surgery (baseline). For six weeks post-surgery, recordings were collected twice weekly at night (6pm-6am) for SCI animals and once weekly for sham and naïve animals.

**Figure 1:**
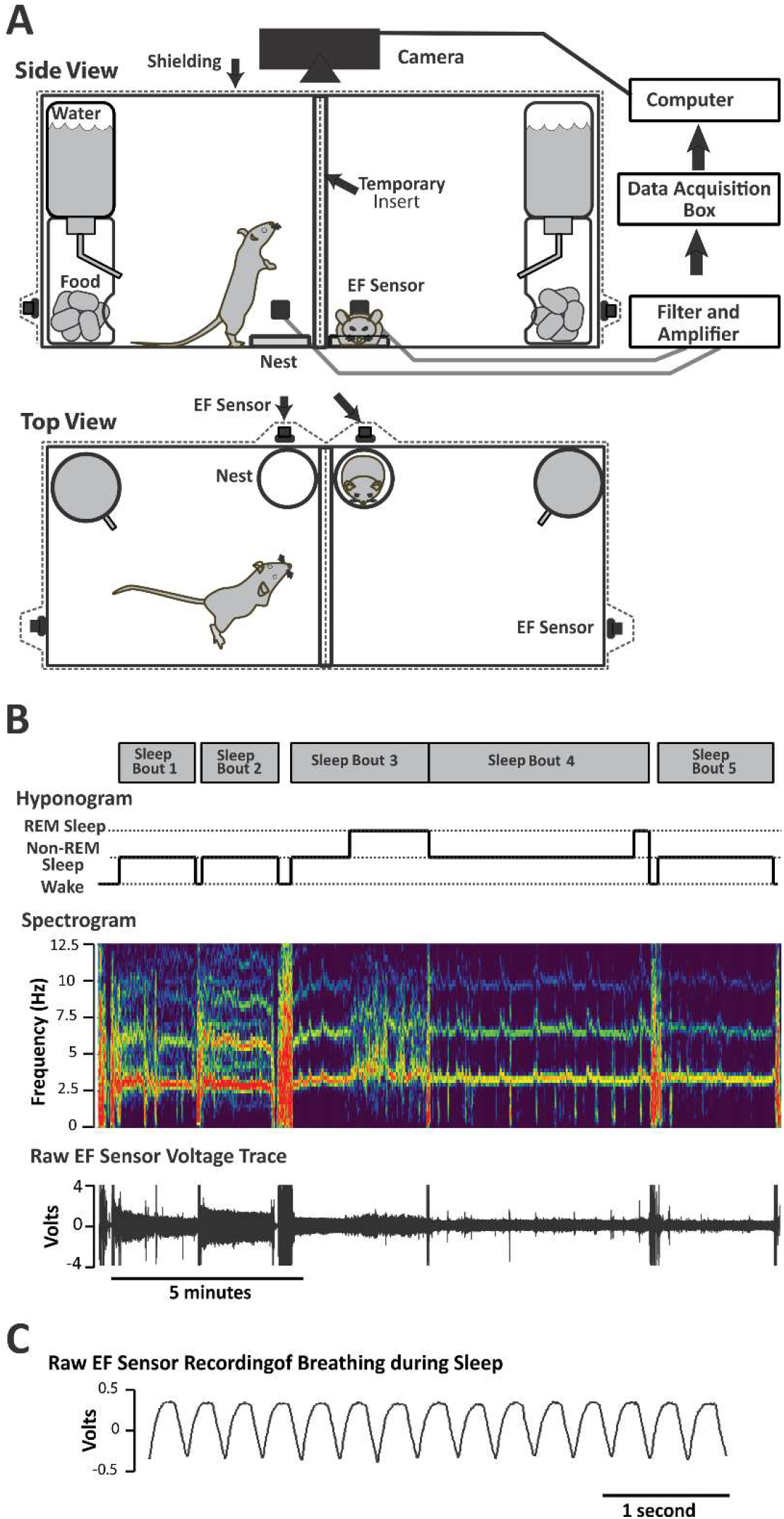
<u>Methodology for non-invasive monitoring of animal sleep and respiration</u>. **A)** Home cage instrumentation for electric field sensing. A total of 4 individual EF sensor were used per cage, with 2 dedicated to each animal. **B)** Representative spectrogram (top panel) and overlaid hypnogram (white trace) obtained from electric field sensor recording (bottom panel) for quantification of sleep events. **C)** Representative recording of breathing with electric field sensor.

### Sleep analysis

Sleep was analyzed from the EF sensor home cage recordings. As described previously ^44, 46^, wake, rapid eye movement (**REM**) sleep, and **non-REM** sleep were quantified through changes in animal movement and respiration (**Figure 1B**). Weekly averages were calculated for total sleep time (**TST**), percent time spent in each arousal state, average sleep bout duration (**SBD**), average REM sleep event duration, REM sleep latency, percentage of sleep bouts containing REM sleep, sleep fragmentation index (**SFI**: total awakenings per hour of sleep), and micro arousal index (**MAI**: total micro arousals < 60 seconds per hour of sleep).

### Respiration analysis

Respiration was detected during non-REM sleep and quiet wake by the EF sensors (**Figure 1C**). Respiratory rate (**RR**) is the frequency at which the animal breathes and RR variability (**RRV**) is the standard deviation of the instantaneous frequencies for each breath during a non-REM event. Weekly values for RR and RRV were calculated using a custom MATLAB script containing 0.1 Hz HP and 10 Hz LP 4^th^ order Butterworth filters.

### Mechanical sensitivity

Mechanical sensitivity was measured using Chaplan’s up-down protocol to calculate 50% paw withdrawal threshold (**PWT**) ^47^. 50% PWT was assessed three times prior to surgery (baseline) and once weekly afterwards for 6 weeks on the same day as overnight physio-behavioral recordings were taken. Single-animal values were obtained by averaging the 50% PWT for both hind paws.

### Isolation of multisegmental DRGs

An isolated spinal cord mouse preparation ^48^ was extended to include DRG from segments C8 through S1. Dorsal and ventral roots were cut to isolate the DRGs and prepared as previously described ^48^. All recordings were undertaken at room temperature.

### Extracellular DRG recordings

Dura and pia were removed around the DRG and glass suction electrodes (200-250 μm tip diameter) were positioned on lumbar (L1-L6) DRGs to record spontaneous afferent population activity for a minimum of 25 seconds (50 kHz, Digidata 1322A 16 Bit DAQ, Molecular Devices, U.S.A., **Figure 2**). Signals were amplified (5000x) and filtered (LP 3 kHz at recording, LP 5Hz post-hoc).

**Figure 2:**
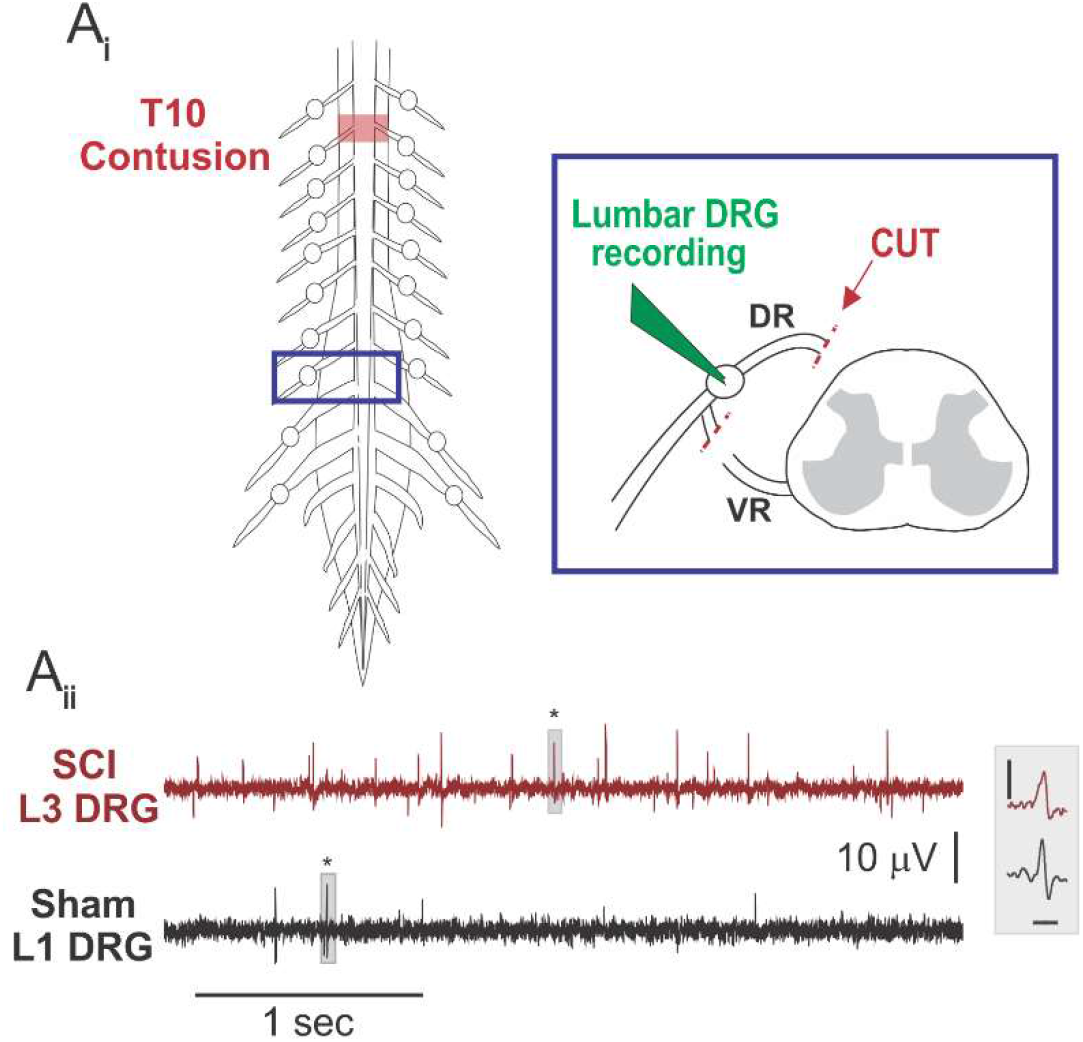
<u>Methodology for electrophysiology for isolated DRG preparation</u>. **A**_**i**_) A schematic of the isolated DRG preparation. Following exposure of the spinal cord and peripheral aspects of primary afferents (DRG, dorsal root, spinal nerve) from C8-S1 segments, the dorsal roots (**DRs**) and ventral roots (**VRs**) were cut at each segment, leaving only a section of the DRs with the attached DRGs and the surrounding ribs and musculature (not shown). **AA**_**ii**_) Example recordings of spontaneous activity in the lumbar (**L**) L3 DRG from the SCI (red) and L1 DRG from the Sham (black) populations. Displayed is a single sweep from maximally active DRGs from 2 different animals. Single units marked with * are expanded in the inset (scale bars are 2 ms and 10 μV).

Spontaneously active **(SA)** and silent DRG spike counts were measured with Clampfit software (v 10.7 Molecular Devices). DRGs were considered SA (**SA DRGs**) with >1 spike within 25 secs. DRG firing frequency was determined by spike frequency within 5 secs bins. Due to the non-uniform sampling of DRGs across animals, a *weighted* SA DRG incidence value was calculated given as the ratio of recorded lumbar DRGs of the 12 is the total possible per animal (#/12).

### Statistical analysis

Statistical differences in behavioral metrics of sleep, respiration, and mechanical sensitivity were determined using 2-Way ANOVA, with a post hoc Sidak’s multiple comparison’s test (α =0.05). Two-tailed Mann-Whitney tests were used to analyze DRG firing frequencies and weighted incidences of SA DRGs. Residual changes from baseline in each animal were quantified for 50% PWT, sleep metrics, and RRV at postoperative week 6 then correlated with post-mortem electrophysiology using Pearson correlation analysis. All statistical analyses were performed using Statistica (Tibco Software Inc. Palo Alto, California) and, unless otherwise stated, all data are presented as mean ± standard deviation.

## RESULTS

### SCI results in chronic hindpaw hypersensitivity

Mechanical sensitivity was assessed to confirm the injury caused mechanical hypersensitivity in agreement with previous findings ^49^. SCI animals developed persistent heightened mechanical sensitivity beginning at week 2 relative to Naïve and Sham groups and SCI baseline (**Figure 3 and Table 1**).

**Figure 3:**
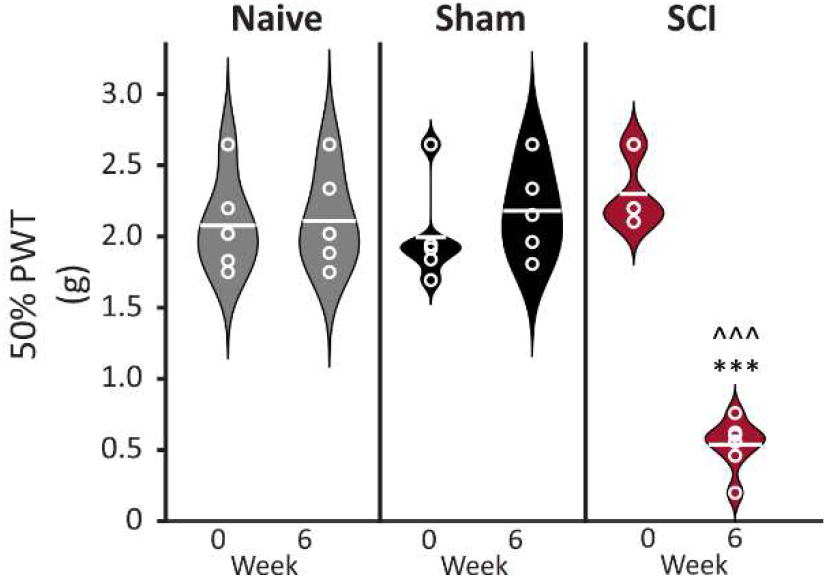
<u>Mechanical Sensitivity</u>. Graph is probability density violin plot of animal averages (circles) with group averages (black and white horizontal lines). Mechanical sensitivity (50% PWT) is heightened after SCI. The p-values are: SCI vs Sham at week 6-*p<0.05, ** p<0.005, *** p<0.0005; baseline vs week 6 in the SCI group-^^^p<0.0001.

**Table 1:**
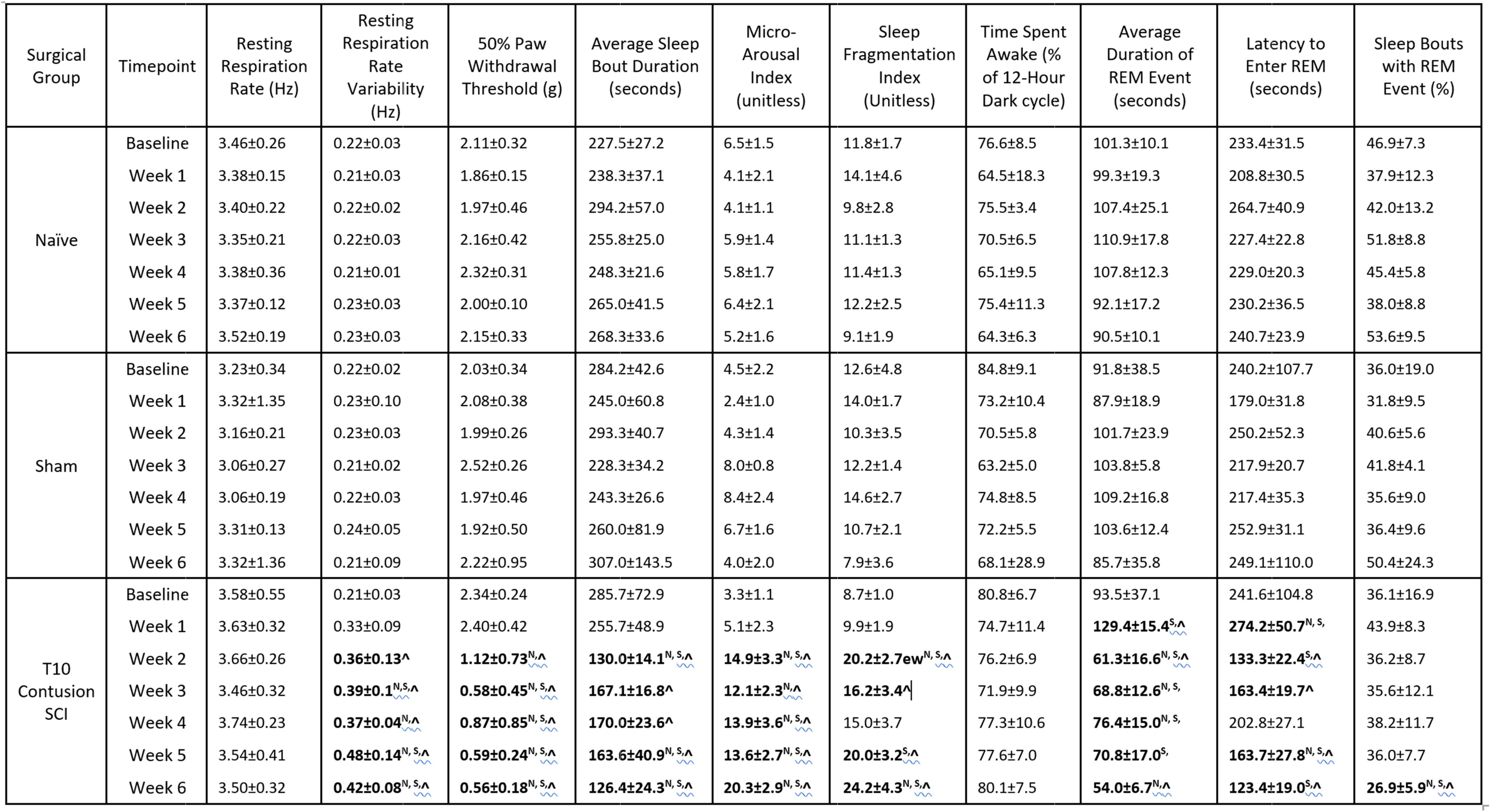
Values are shown as mean ± STD with statistically significant values bolded. ^**N**^: Difference from Naïve at respective timepoint (p<0.05). ^**S**^: Difference from Sham at respective timepoint (p<0.05). **^**: Difference from baseline of respective group (p<0.05). Resting respiration rate (RR) and percentage of time spent awake were not affected by SCI. By 6 weeks post-injury, SCI animal mechanical sensitivity (50% PWT), sleep fragmentation indices, and respiration rate variability (RRV) were significantly different from SCI baseline and 6 week Naïve and Sham group values. Specifically, RRV, 50% PWT, sleep bout duration, sleep fragmentation index (SFI), and micro-arousal index (MAI) showed progressive changes starting at week 2 post-injury, while changes in REM sleep did not solidify until week 6 post-injury.

### Respiration becomes more erratic after SCI

The average resting RR did not show significant differences between treatment groups or across time (**Figure 4 and Table 1**). At baseline, all treatment groups exhibited a consistent respiratory rate variability (RRV) averaging 0.21±0.02 Hz. However, RRV increased by week 2 for the SCI group relative to SCI baseline and increased relative to Naïve and Sham groups at all time periods after two weeks.

**Figure 4:**
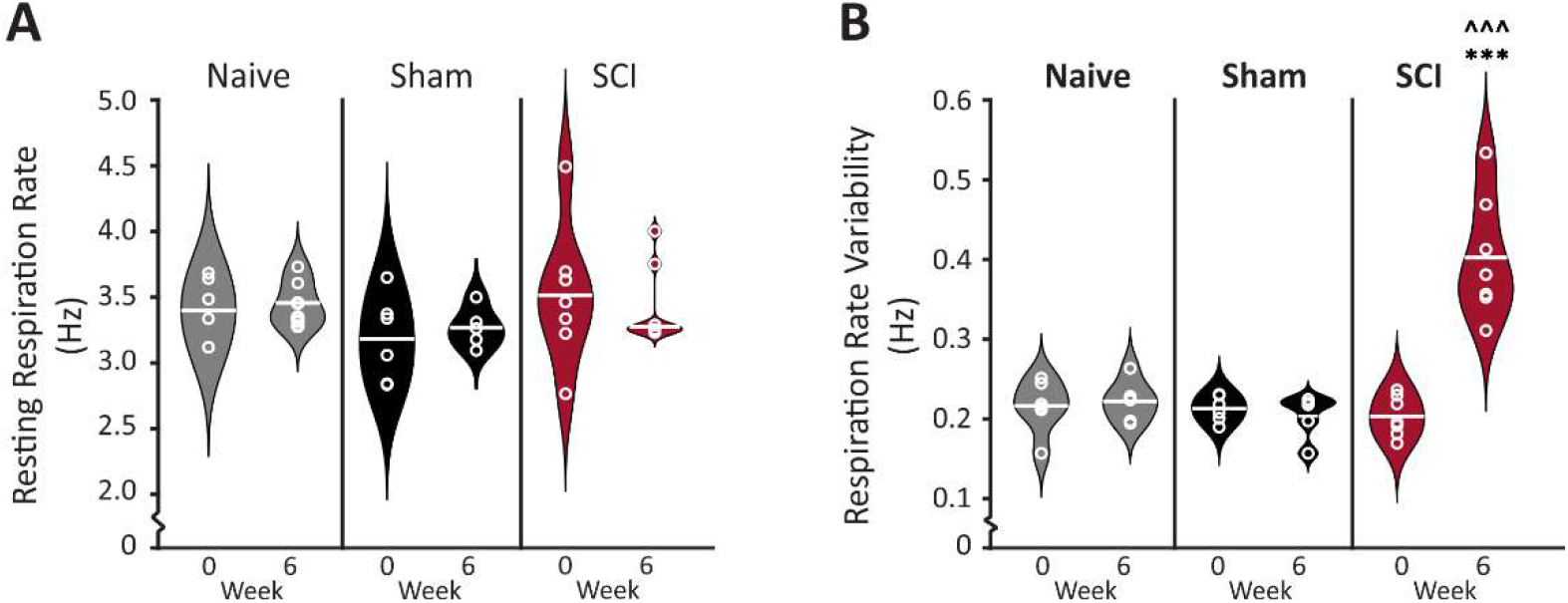
<u>Respiration Changes After SCI</u>. Graphs are probability density violin plots of animal averages (circles) with group averages (black and white horizontal lines). A) Average respiratory rate appears unaffected by SCI, while B) respiratory rate variability differs significantly when comparing SCI vs Sham at postoperative week 6; and baseline vs week 6 in the SCI group. The p-values are: SCI vs Sham at week 6-*p<0.05, ** p<0.005, *** p<0.0005; baseline vs week 6 in the SCI group-^^^p<0.0001.

### Sleep becomes fragmented after SCI

Total sleep time was not significantly different between treatments or across time (**Figure 5 and Table1**). Beginning at week 2, SCI animals developed progressive fragmented sleep as measured by decreased sleep bout duration, increased SFI, and increased MAI relative to Naïve and Sham groups and SCI baseline. In SCI animals, REM sleep duration, REM latency, and the percentage of sleep bouts containing REM sleep decreased at later timepoints relative to Naïve and Sham groups and SCI baseline. Specifically, changes in REM sleep developed later than both mechanical sensitivity and initial sleep fragmentation.

**Figure 5:**
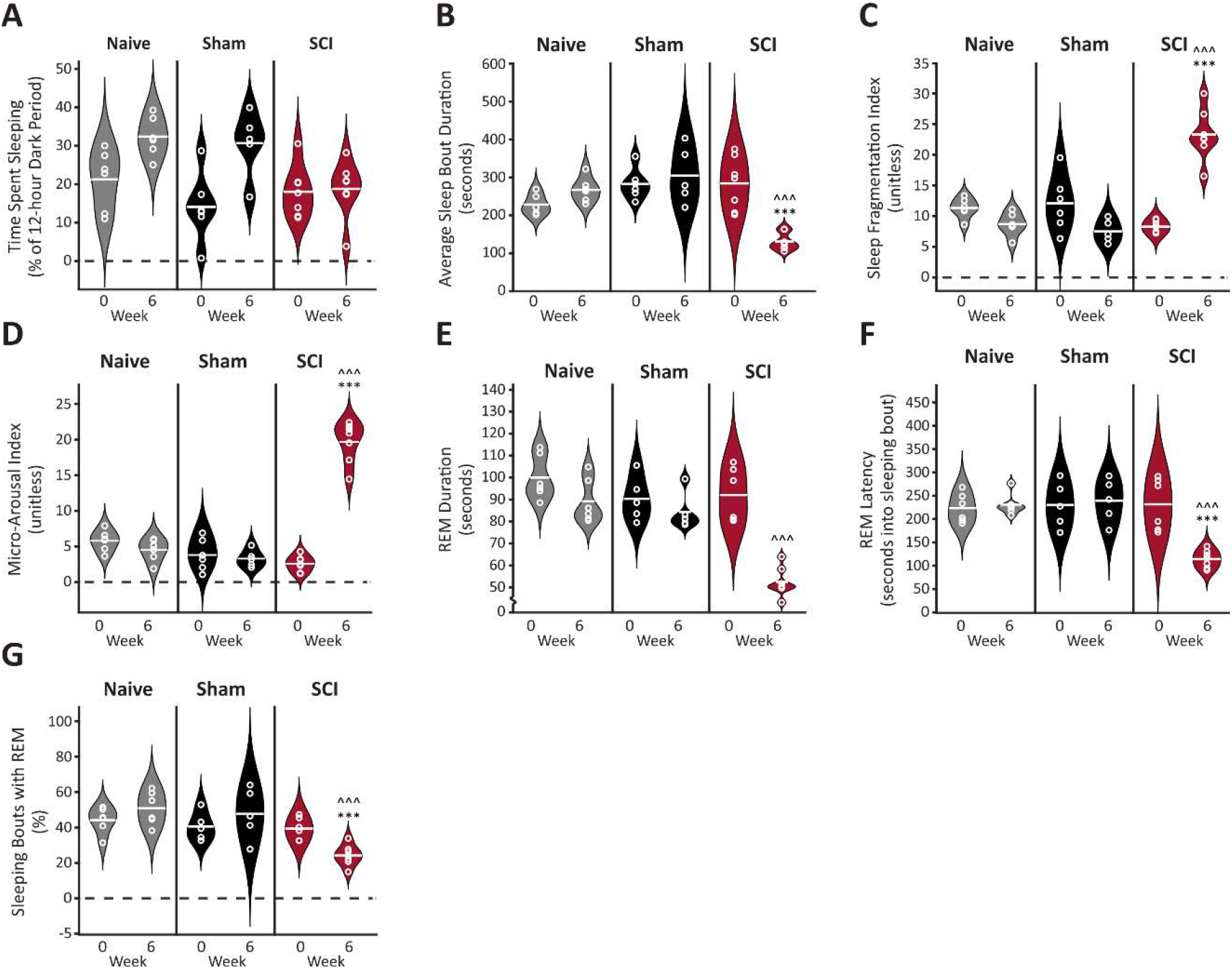
<u>Sleep Changes After SCI</u>. Graphs are probability density violin plots of animal averages (circles) with group averages (black and white horizontal lines). A) While time spent sleeping was unaffected by SCI, B) average sleep bout duration, C) SFI, and D) MAI during a 12-hour dark cycle were significantly different when comparing SCI vs Sham at postoperative week 6; and baseline vs week 6 in the SCI group. E) REM sleep duration, F) REM sleep latency, and G) the percentage of sleep bouts containing REM sleep were all reduced in SCI animals at postoperative week 6 relative to their baseline and sham animals at postoperative week 6 (except for REM sleep duration) suggesting premature arousal to wake, an increase in sleep pressure, and an overall reduction in sleep quality. The p-values are: SCI vs Sham at week 6-*p<0.05, ** p<0.005, *** p<0.0005; baseline vs week 6 in the SCI group-^^^p<0.0001.

### Greater spontaneous nerve activity emerges after SCI

To assess below-level spontaneous primary afferent activity, spontaneous nerve population activity in L1-L6 DRGs was recorded bilaterally from SCI and Sham animals 6 weeks after injury. Compared to Sham (0.6 ± 0.3 Hz), SCI animals exhibited greater spontaneous activity (2.5 ± 1.2 Hz), as characterized by the maximal DRG firing frequency (p < 0.005, **Figure 6**). The weighted incidence of SA DRGs in the SCI population (29.8 ± 16.5%) was significantly greater than Sham (12.5 ± 4.6%, p < 0.5, **Figure 6**).

**Figure 6:**
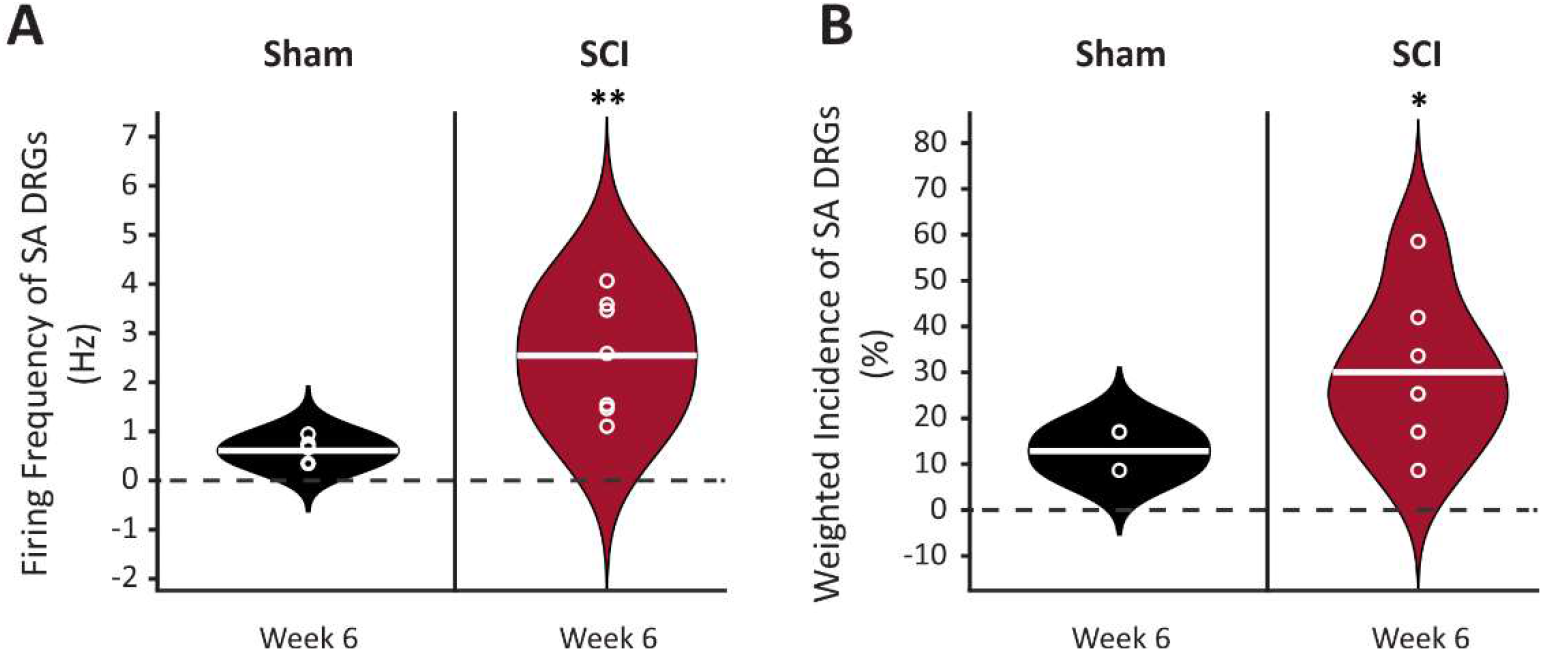
<u>Spontaneous Afferent Firing</u>. Graphs are probability density violin plots of animal averages (circles) with group averages (black and white horizontal lines). A) The maximal observed firing frequency and B) weighted incidence of SA lumbar DRGs in the SCI group are significantly greater than that of the Sham group. SCI (n=7), Sham (n=5). The p-values are: SCI vs Sham at week 6-*p<0.05, ** p<0.005, *** p<0.0005; baseline vs week 6 in the SCI group-^^^p<0.0001.

### Spontaneous Hyperexcitability Correlates with Changes in Evoked Hyperexcitability, Respiration, and Sleep Changes after SCI

Correlative relationships between behavioral changes and spontaneous afferent activity were identified (**Figure 7**). Though direct comparisons between behavior and electrophysiology yield no clear correlations, residualized behavior measures between postoperative week 6 and baseline (i.e. the change measured at week 6 from each animal’s respective baseline) reveal significant relationships: the change in mechanical sensitivity (50%PWT) negatively correlates with the weighted incidence of SA DRGs, the change in SFI positively correlates with the weighted incidence of SA DRGs, and the change in RRV positively correlates with DRG firing frequency. These findings link sleep and respiratory features to changes in spontaneous afferent activity and neuropathic pain after SCI.

**Figure 7:**
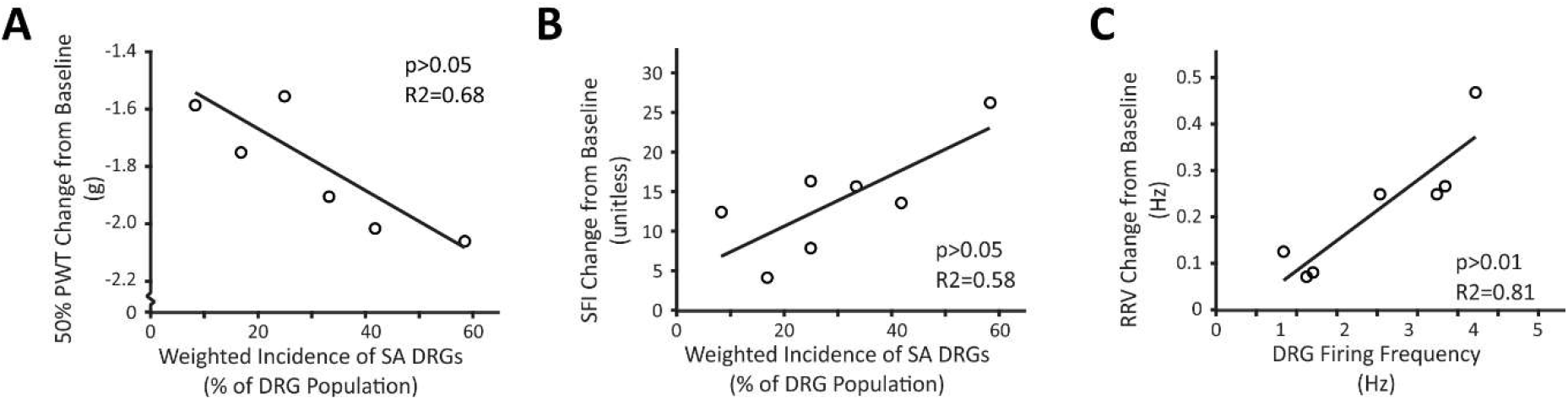
<u>Spontaneous hyperexcitability metrics correlate with changes in evoked hyperexcitability, respiration, and sleep after SCI</u>. A) Incidence of SA DRGs in the SCI group has a negative correlation with the change in 50% PWT threshold (compared to baseline), measured at postoperative week 6. B) In the SCI group, the change in the SFI, which captures the number of sleep disruptions per hour, at week 6 (compared to baseline) has a positive correlation with the incidence of below-injury-level SA DRGs. C) Maximal firing frequency has a positive correlation with the change in respiration rate variability (compared to baseline) at week 6. Linear associations between metrics were determined by Pearson correlation and regression analysis. Each data point represents a single animal.

## DISCUSSION

This work links spontaneous afferent activity, behavioral mechanosensitivity measures consistent with neuropathic pain to measures of sleep and respiration in naturally behaving mice. While peripheral hyperexcitability has been previously observed in SCI models of neuropathic pain, here we found correlations between spontaneous lumbar DRG activity (incidence and firing frequency) and metrics of sleep and respiratory dysfunction – complications frequently observed in SCI patients. The findings suggest that in addition to examining primary afferent activity, non-contact recording of sleep and respiratory features may serve as a useful tool for identifying the emergence of persistent spontaneous neuropathic pain in animal models of SCI. Below we discuss results of SCI -induced changes in sleep, respiration and sensory hyperexcitability changes in separate sections prior to providing our overall perspective.

### Sleep changes after SCI

Currently there is very limited preclinical animal studies devoted to assessing sleep changes after SCI. Only one study looked at and observed sleep dysfunction independent of respiration^50^. This study was undertaken in rat and follow changes for 15 days after complete lower thoracic spinal tansection and observed a transient reduction in sleep efficiency during the light cycle.

The technological advance of the EF sensors has enabled noninvasive quantification of sleep changes in rodents after SCI ^44^. Though the time spent asleep during the 12-hour dark cycle (6pm-6am) was unchanged, SCI animals had more fragmented and lower quality sleep relative to their sham and naïve counterparts. There were meaningful reductions in average sleep bout duration (SBD), REM sleep duration, REM sleep latency, and the percentage of sleep bouts containing REM sleep. Sleep fragmentation index (SFI) representing total number of awakenings/hr of sleep, and micro arousal index (MAI) representing total number of <60 seconds micro arousals/hr of sleep were increased after SCI. The increased brief arousals and reduced duration of sleep bouts caused compensatory increased number of sleep events after SCI, presumably reflecting experienced increased sleep pressure.

Since REM sleep occurs at the end of a sleep cycle, it is intuitive to link these observed reductions in REM sleep to reduced SBD associated with observed increased sleep fragmentation and micro arousals. If sleep events are shortened and without observed REM sleep, it is possible the animal did not achieve normal transitions for restorative sleep. Factors responsible for sleep compromise may include increased pain pathway recruitment of sympathetic drive, given known correspondence with mechanical hypersensitivity and in other studies showing evidence of ongoing spontaneous pain ^6, 51^. Regardless of mechanism, sleep dysfunction would contribute to overall stress burden after SCI and, through actions on common stress axis pathways contribute to overall disease burden in SCI.

### Respiration changes after SCI

While cervical SCI clearly impair respiration due to direct injury to respiratory neural circuits that impact sleep ^52–54^, there is little consideration on whether SCI -induced neural dysfunction in other areas may interact with an impact respiration. Nonetheless, recent work has shown that, when recorded in restraint cylinders, respiratory rate increases after thoracic contusion or hemisection injury that may precede to correspond with the emergence of allodynia ^40, 41^. In comparison, and recordings of respiratory rate in naturally behaving mice, we observed no change in average respiration rate but with increases respiratory rate variability (RRV). These changes also emerged coincident with measures of mechanical hypersensitivity.

### Neural basis of neuropathic pain after SCI

In SCI animals, we showed a correlation between the change in paw withdrawal threshold (PWT) and the and dorsal ganglia (DRG) afferent firing incidence, where a greater reduction in PWT was observed in animals with a higher incidence of spontaneously active DRGs. DRG firing frequency and incidence were not correlated, suggesting they may capture disparate aspects of ongoing pain or an alternate non-pain encoding sensory signaling pathway (e.g. metabolic status, baroreceptors, hypoxia). The incidence of spontaneously active DRGs may determine the size/number of dermatomal regions impacted by neuropathic pain—where a larger incidence could be associated with a larger region impacted. Firing frequency may encode the intensity of pain experienced by a single region which could explain the link between PWT and DRG incidence as the rodent hindpaw is innervated by axons with cell bodies in L3-L6 DRGs ^55^. Broader hyperexcitability of these lumbar primary afferents may contribute to greater hindpaw sensitivity.

Spontaneous primary afferent activity has been identified as a driver of chronic pain in clinical ^56–58^ and preclinical ^6, 7, 18, 19, 22^ studies. Key studies conducted in rodent contusion models of SCI revealed potential links between stimulus-evoked hypersensitivity and spontaneous firing of primary afferents below the level of injury ^6, 16, 22, 23^. Here, we recorded activity from intact DRGs across multiple vertebral segments to confirm development of increased spontaneous afferent activity after SCI. Though our recording technique did not permit resolution to identify the class of primary afferent nerves within the signal population, there is evidence suggesting spontaneous nerve activity after SCI includes signals from A and C fibers which have been associated with neuropathic pain including in the T10 contusion SCI animal model ^6, 7, 15, 16, 18, 19, 22, 58, 59^.

## CONCLUSIONS

We show that a commonly used SCI animal model of neuropathic pain is also associated with co-emergent sleep dysfunction and increased respiratory rate variability, thereby providing a broader understanding of the magnitude of overall stress burden initiated by neural circuit dysfunction after SCI. Further studies are warranted to better understand the nature of their interactions be they cause-effect relations or independent parallel injury-induced phenomenon on distinct pathways impacted by SCI as such insights may inform implementation of therapeutic control strategies after SCI.

## ACKNOWLEDGMENTS

We thank Mallika Halder for assisting with the animal surgeries; Sandra Garraway and Karmarcha Martin for training us in the development of the T10 contusion model and permitting the use of the Infinite Horizon impactor.

## RESEARCH CONTRIBUTIONS

Animal surgeries were performed by Shaquia Idlett-Ali and Mallika Halder. Tissue isolation, electrophysiology experimentation, and analysis was performed by Shaquia Idlett-Ali. Behavioral testing and analysis was conducted by Heidi Kloefkorn. Shaquia Idlett-Ali, Heidi Kloefkorn and Shawn Hochman were involved in preparation of the manuscript.

